# AFilter: Improved Antibody Epitope Prediction by Machine Learning-Optimized Interface Energy Filtering of AlphaFold3-Predicted Complex

**DOI:** 10.64898/2026.08.16.745116

**Authors:** Xiaoyu Liu, Yu YoSean Wang

## Abstract

AlphaFold3 (AF3) predicts protein-complex structures from sequence with near-experimental accuracy on many targets, substantially lowering the cost of mechanistic and therapeutic discovery. However, application to antibody epitope prediction is hampered by an approximately 63% failure rate. Comparing successful and failed AF3 predictions across antibody-antigen and nanobody-antigen complexes, we found that failed predictions share a distinctive energetic signature: distorted CDR-loop geometries and elevated van der Waals strain at the interface. Building upon these observations, we developed a machine learning-based interface energy filtering framework, designated AFilter, capable of eliminating over 90% of erroneous predictions while retaining >90% of true positives. Compared with ipTM-based filtering, AFilter improved accuracy from 82.7% to 97.7% for nanobody-antigen complexes and from 79.4% to 96.3% for antibody-antigen complexes, while simultaneously raising the true positive rate from 69.8% to 96.4% and from 63.1% to 92.5%, respectively. When applied to NeuroMab antibodies of unknown structure, AFilter prioritized high-confidence epitope predictions that AF3 sampling alone could not reliably surface. As a lightweight post-hoc filter (<5% computational overhead) that requires no re-docking, AFilter is directly compatible with existing AF3 prediction pipelines and, in principle, transferable to other diffusion-based complex predictors, providing a practical quality-assurance layer for antibody epitope mapping in early-stage drug discovery.

## Introduction

The epitope — the surface region of an antigen recognized by an antibody’s paratope — dictates the binding affinity, specificity, and functional outcome of the antibody-antigen interaction [1], and identifying it is therefore a prerequisite for antibody drug development. Anti-PD-1 antibodies such as pembrolizumab and nivolumab, for example, occupy epitopes on PD-1 that sterically occlude PD-L1 engagement [2, 3], and broadly neutralizing antibodies against SARS-CoV-2 RBD achieve cross-variant protection by targeting epitopes of low mutational escape [4, 5]. Epitope configuration also directly influences target druggability and antibody developability [6], informing downstream antibody engineering and clinical success [7, 8].

Experimental epitope mapping, however, occurs late in the antibody pipeline and constitutes a major bottleneck [9, 10]. High-resolution methods — X-ray co-crystallography and cryo-EM — resolve antibody-antigen contacts at near-atomic resolution but require purified protein and weeks to months of effort per target [11]. Faster alternatives such as hydrogen-deuterium exchange mass spectrometry (HDX-MS) or alanine/shotgun mutagenesis with SPR/BLI readouts either suffer from artifacts or demand iterative construct cycles [12, 13]. Modern antibody campaigns routinely generate hundreds to thousands of clones [14], so epitope characterization frequently dictates the rate at which lead candidates can be triaged.

Deep-learning structure prediction has opened a path to computational epitope mapping [15], but AlphaFold-Multimer (AFM) reaches acceptable docking quality on only ∼20–30% of antibody-antigen pairs [16]. AlphaFold3 (AF3) replaces the Evoformer with a Pairformer module and introduces a diffusion-based structure generator, broadening conformational sampling in a single forward pass [17]. Hitawala and Gray reported high-accuracy AF3 docking rates of ∼9–13% for antibody- and nanobody-antigen pairs under single-seed sampling, and the AF3 authors noted that scaling to 1,000 seeds raises the overall success rate to ∼60% [17, 18]; under the more practical three-seed protocol used here, only ∼37% of predictions reach medium or higher quality (DockQ > 0.49; Figure 1C of this work). Several structural features of antibody-antigen interfaces contribute to this persistent failure rate. CDR loops protrude into concave epitope surfaces and demand precise shape complementarity unlike the flatter interfaces of typical protein-protein complexes [19], and CDR H3 in particular varies in length from 2 to >30 residues with diverse conformations, posing a sampling challenge that AF3’s limited diffusion trajectory may not fully address [20, 21]. Taken together, these observations indicate that raw AF3 outputs are not directly usable for antibody-epitope prediction and motivate a post-prediction filter that separates correct from incorrect docking poses.

**Figure 1.**
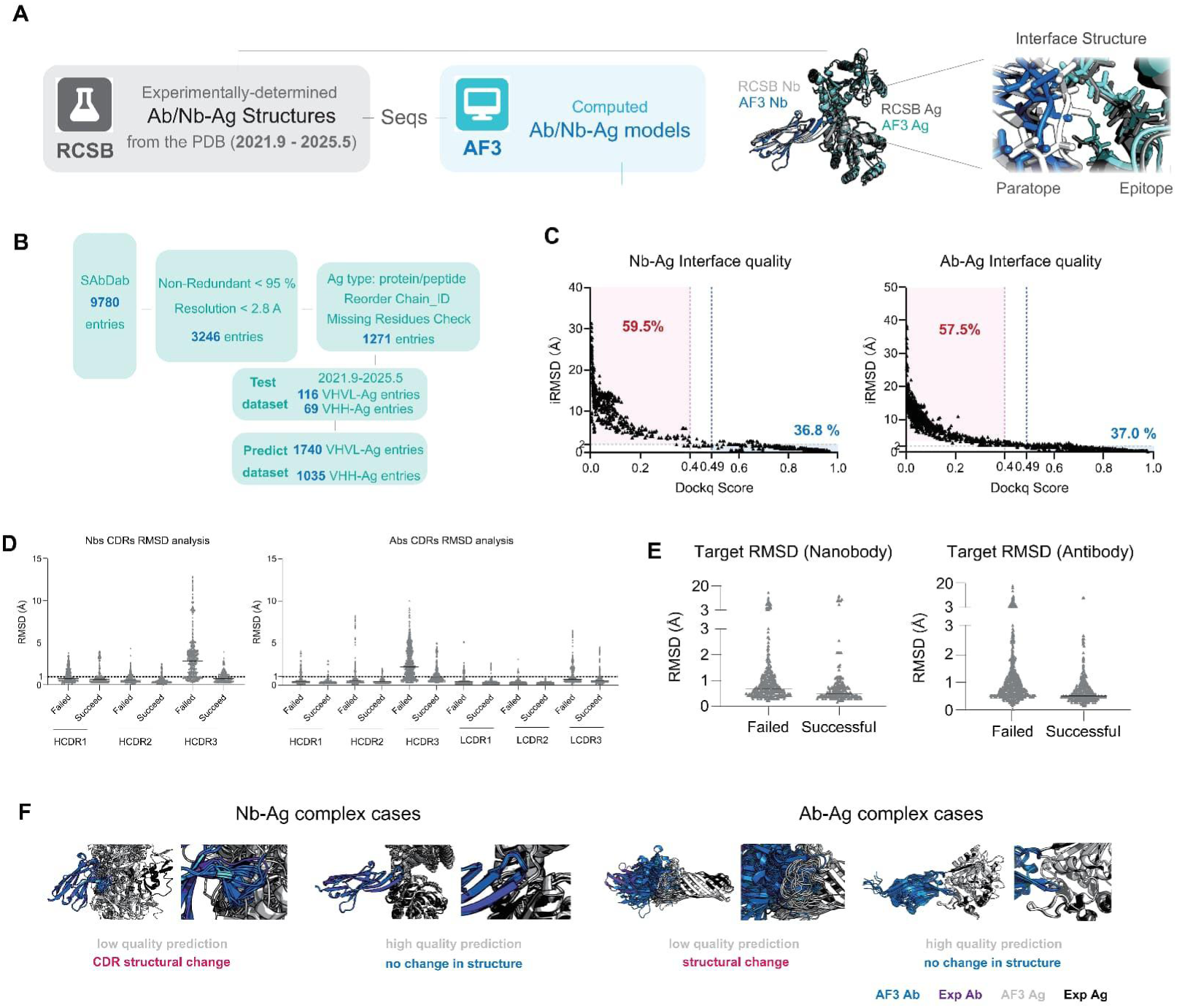
Conformational Changes at Antibody-Antigen Interfaces in Failed AF3 Predictions. (A) Overview of the benchmarking pipeline. Experimentally determined antibody/nanobody-antigen complex structures deposited in the PDB between September 2021 and May 2025 were retrieved, and their sequences were submitted to AlphaFold3 to generate predicted complex models. Predicted and experimental interface structures were then compared at the paratope-epitope level. (B) Data curation workflow. Starting from 9,780 SAbDab entries, sequential filters for non-redundancy (<95% antibody sequence identity), resolution (<2.8 Å), antigen type (protein/peptide), chain-order consistency, and missing-residue completeness yielded 1,271 entries for the reference distribution, 116 VHVL-Ag and 69 VHH-Ag entries for the test dataset, and 1,740 VHVL-Ag and 1035 VHH-Ag predicted conformations. (C) AF3 docking quality assessed by DockQ score versus interface RMSD (iRMSD) for nanobody-antigen (left) and antibody-antigen (right) complexes. Pink-shaded regions indicate failed predictions; percentages denote the proportion of predictions in each quality regime. Dashed lines mark the DockQ = 0.49 and DockQ = 0.40 thresholds. (D) CDR loop RMSD relative to experimental structures for successful versus failed AF3 predictions, stratified by CDR region. Left: nanobody CDRs (HCDR1, HCDR2, HCDR3); right: antibody CDRs (HCDR1, HCDR2, HCDR3, LCDR1, LCDR2, LCDR3). CDR H3 exhibits the largest deviations in failed predictions. (E) Antigen-side interface RMSD (Target RMSD) for successful versus failed predictions in nanobody-antigen (left) and antibody-antigen (right) complexes. Deviations are smaller than those observed on the antibody/nanobody side. (F) Representative structural overlays of AF3-predicted (blue) and experimental (gray) complexes for nanobody-antigen (left two panels) and antibody-antigen (right two panels) cases, illustrating CDR structural distortions in low-quality predictions and preserved CDR conformations in high-quality predictions.

To address this need, we assembled and analyzed a curated dataset of 1,509 nanobody-antigen and 2,722 antibody-antigen complexes derived from the SAbDab database [22, 23] and corresponding AF3 predictions. Systematic comparison of successful and failed predictions revealed that incorrectly docked complexes exhibit distinctive abnormalities readily quantifiable by physics-based energy evaluation: specifically, elevated van der Waals repulsion (fa_rep), distorted backbone dihedral geometries (rama_prepro), and unfavorable electrostatic profiles at the interaction interface, as scored by the Rosetta REF2015 all-atom energy function [24]. These observations are consistent with the general principle that native-like protein-protein interfaces are characterized by tight atomic packing and favorable energetics [19], and align with recent work demonstrating that physics-based rescoring can improve the discrimination of near-native docking poses generated by deep learning methods [25, 26]. Building on these findings, we developed AFilter, a machine learning classifier trained on Rosetta-derived interface energy features to distinguish high-quality from low-quality AF3 predictions. On held-out validation data, AFilter eliminated over 90% of false-positive predictions while retaining approximately 90% of true positives, and maintained comparable performance on an independent test set of complexes determined after the training data cutoff. We further demonstrate AFilter’s practical utility by applying it to map previously uncharacterized epitopes of NeuroMab antibodies. Collectively, our results show that integrating physics-based interface energy evaluation with deep learning-generated structural models provides a robust and computationally efficient strategy for improving the reliability of antibody epitope prediction.

## Materials and Methods

### Data Collection

Antibody-antigen and nanobody-antigen complex structures, together with their corresponding FASTA sequences, were retrieved from the Structural Antibody Database (SAbDab) [22, 23], covering accession dates from January 1, 1970 to May 7, 2025. To ensure structural reliability and reduce redundancy, only complexes resolved at <2.8 Å and sharing <95% antibody sequence identity were retained. A temporal subset of these complexes — those with PDB accession dates after September 30, 2021 (the AF3 training data cutoff [17]) through May 2025 — was used to generate AF3 prediction models via the AF3 server. For each complex, three random seeds were employed, with five independent diffusion sampling runs per seed, yielding 15 predicted conformations per antibody-antigen pair. This set of predicted structures served two purposes: (1) benchmarking AF3’s predictive accuracy on antibody-antigen docking, and (2) providing labeled training and validation data, alongside the full SAbDab experimental structure collection, for a machine learning model designed to assess AF3 prediction quality. Specifically, the temporally held-out subset comprised 116 antibody-antigen and 69 nanobody-antigen pairs and was used both for AF3 benchmarking and for ML training/validation; the remaining SAbDab entries deposited on or before 30 September 2021 (1,509 nanobody-antigen and 2,722 antibody-antigen complexes in total) were used only as the experimental reference distribution for energy-feature characterization and were not included in AF3 prediction or model training.

### Structural Data Preprocessing

All experimental and AF3-predicted structures were processed using PyRosetta [27] to remove non-amino-acid components (ligands, water molecules, and ions) and to renumber residue identifiers sequentially. The chain order of AF3-predicted structures was aligned with that of the corresponding experimental PDB files to enable consistent computation of interface RMSD (iRMSD) [28] and DockQ scores [29]. Because experimental structures frequently contain missing heavy atoms and non-standard hydrogen placement that can distort energy calculations, all hydrogen atoms were first stripped using PyMOL [30] and then rebuilt with PyRosetta’s hydrogen-packing routines to ensure uniform protonation states across all structures.

### Interface Energy Analysis

Antibody and antigen chains within each complex were annotated using PyIgClassify [31]. Interface contact residues were defined as those with any inter-chain heavy-atom distance <6 Å, identified using pdb2sql [32], and partitioned into antibody-side and antigen-side subsets. For each contact residue, 18 per-residue energy terms from the Rosetta REF2015 all-atom energy function [24] were extracted using PyRosetta (expressed in Rosetta Energy Units, REU). These terms include van der Waals attraction (fa_atr) and repulsion (fa_rep), Coulombic electrostatics (fa_elec), solvation (fa_sol), backbone dihedral preferences (rama_prepro), hydrogen bonding (hbond_sr_bb, hbond_lr_bb, hbond_bb_sc, hbond_sc), side-chain rotamer energy (fa_dun), and others. The 18 antibody-side and 18 antigen-side mean per-residue energies were concatenated to yield a 36-dimensional feature vector for each complex.

### Machine Learning-Based Filter of AF3-Predicted Complex Quality

We treated AF3 quality assessment as a binary classification problem over the 36-dimensional interface-energy feature vectors. Ground-truth labels were assigned using joint DockQ and iRMSD criteria: predictions with DockQ > 0.49 and iRMSD < 2.0 Å were labeled as correct (positive), and those with DockQ ≤ 0.40 or iRMSD ≥ 2.0 Å were labeled as incorrect (negative); predictions falling between these thresholds were excluded to reduce label ambiguity. The labeled dataset was split into training (80%) and validation (20%) subsets with stratified sampling. We note that the descriptive comparisons of CDR-loop RMSD, interface RMSD, and per-residue fa_rep distributions presented in Figures 1D–E and 2E use the simpler binary contrast DockQ > 0.49 versus DockQ ≤ 0.49 across all 15 predicted conformations per pair, whereas model training and validation rely on the stricter labeling scheme above with the ambiguity gap excluded.

Five classification algorithms-Random Forest [33], Gradient Boosting Decision Tree (GBDT) [34], AdaBoost, Decision Tree, and Naive Bayes — were trained across 100 random seeds to assess stability. Feature selection was performed by ranking features using mean decrease in impurity from the Random Forest classifier, averaged over 500 independent runs, and the top 10 features were retained for final model training. Each selected model was retrained using five-fold cross-validation. Model performance was compared on the basis of area under the receiver operating characteristic curve (AUC-ROC). Random Forest was selected as the final classifier owing to its consistently high AUC and superior robustness to outliers at the current dataset scale (1,000–3,000 samples). The classification threshold for each model was adjusted to maintain a true positive rate ≥ 90%.

For independent evaluation, newly released antibody-antigen and nanobody-antigen complexes (PDB accession dates: June-November 2025), filtered by the same resolution and redundancy criteria, were used as an external test set. AF3 predictions were generated under the same three-seed, five-sample protocol (15 conformations per pair). Ground-truth labels for the test set followed the same DockQ/iRMSD criteria described above.

### Computational Resources

All structure preprocessing and Rosetta energy calculations were performed on AMD EPYC 7742 processors. AF3 predictions were generated using NVIDIA RTX 2080 Ti and RTX 3090 GPUs. Machine learning model training and inference were carried out using scikit-learn [35] in Python 3.9.

## Results

### 1. Widespread Conformational Changes in Failed AF3-Predicted Epitopes

To benchmark AF3’s epitope prediction accuracy on data unseen during model training, we curated 116 antibody-antigen and 69 nanobody-antigen complexes from SAbDab, applying the temporal, resolution, and redundancy filters described in Methods (Figure 1A, B). AF3 predictions were generated with three random seeds and five diffusion samples per seed, yielding 1,740 antibody and 1,035 nanobody predicted conformations in total. Using stringent quality thresholds — predictions with DockQ > 0.49 and iRMSD < 2.0 Å were labeled as correct (positive), and those with DockQ ≤ 0.40 or iRMSD ≥ 2.0 Å were labeled as incorrect (negative); predictions falling between these thresholds were excluded to reduce label ambiguity — both antibody and nanobody epitope predictions achieved an overall success rate of approximately 36-37% (Figure 1C). When examined at the level of individual antibody-antigen pairs, however, prediction success showed substantial heterogeneity: some pairs were correctly predicted in the majority of seeds, while others failed consistently across all 15 conformations. This pair-to-pair variability may partly reflect the limited sampling depth of 15 conformations per pair, and is consistent with the AF3 authors’ observation that expanding sampling to 1,000 seeds raises the aggregate success rate to approximately 60% [17].

To investigate the structural basis of prediction failure, we compared interface geometries between successful and failed groups. On the antibody side, failed predictions exhibited markedly elevated CDR loop RMSD relative to the corresponding experimental structures, with CDR H3 showing the largest deviations (Figure 1D; p < 0.001, Wilcoxon rank-sum test). This pattern was observed for both conventional antibodies and nanobodies. A similar trend was present on the antigen side of the interface, although the magnitude of the deviations was smaller (Figure 1E), likely because antigen interface regions contain a higher proportion of rigid secondary structure elements compared with the inherently flexible CDR loops on the antibody side. Representative structural overlays illustrate this pattern: in failed predictions, CDR loops — particularly CDR H3 — adopt markedly distorted conformations relative to the experimental structure, whereas in successful predictions the CDR geometry is well preserved (Figure 1F). These results indicate that when AF3 docks an antibody at an incorrect epitope, it compensates by distorting CDR loop conformations — particularly CDR H3 — to force a superficially plausible interface at a non-native binding site. The resulting structural distortions, while potentially satisfying AF3’s internal confidence metrics, introduce non-physical features that can be quantified through per-residue interface energy analysis, as explored in the next section.

### 2. Pervasive van der Waals Over-Repulsion in Failed AF3-Predicted Epitopes

Having established that failed AF3 predictions are accompanied by CDR conformational distortions, we next asked whether they leave a detectable energetic signature. Using the 36-dimensional Rosetta interface energy feature vectors described in Methods, we first assessed feature importance by training Random Forest classifiers across 500 independent runs with varying random seeds and ranking features by their mean decrease in impurity (Gini importance) (Figure 2B). Van der Waals repulsion (fa_rep) on both the antibody side (Ab_fa_rep) and antigen side (Ag_fa_rep) consistently emerged as the two most discriminative features, followed by backbone dihedral preferences (rama_prepro) and electrostatic terms (fa_elec) (Figure 2B).

**Figure 2.**
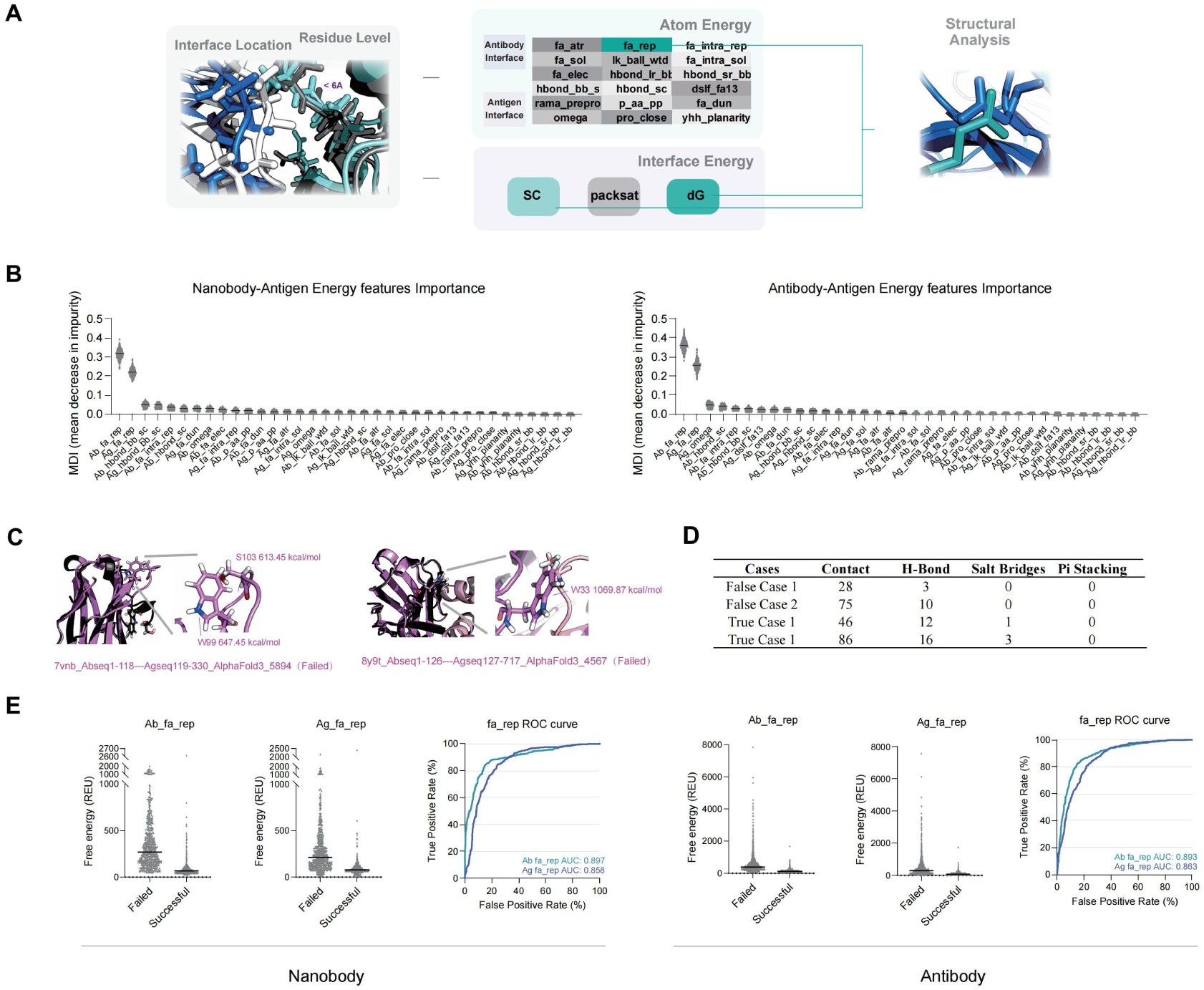
Elevated van der Waals Repulsion at Interfaces of Failed AF3 Predictions. (A) Schematic of the per-residue energy feature extraction pipeline. Interface residues (inter-chain heavy-atom distance <6 Å) are identified, and 18 Rosetta REF2015 energy terms are extracted separately for antibody-side and antigen-side contact residues, yielding a 36-dimensional feature vector per complex. Energy terms are grouped into categories including conformational (rama_prepro), pairwise (fa_rep, fa_atr, fa_elec), solvation (fa_sol), and interface-level metrics (shape complementarity, packstat, dG). (B) Feature importance rankings based on mean decrease in impurity from 500 independent Random Forest runs. Left: nanobody-antigen dataset; right: antibody-antigen dataset. Ab_fa_rep and Ag_fa_rep consistently emerge as the two most discriminative features. (C) Representative structural examples of failed AF3 predictions highlighting specific residues with extreme van der Waals repulsion energies (fa_rep values annotated in REU). (D) Summary table comparing interface interaction properties (contacts, hydrogen bonds, salt bridges, pi stacking) between representative false-case and true-case predictions. (E) Left and center panels: violin plots comparing mean per-residue fa_rep values at the interface for failed versus successful predictions on the antibody/nanobody side (Ab_fa_rep) and antigen side (Ag_fa_rep), shown separately for nanobody-antigen (left group) and antibody-antigen (right group) complexes. Right panels: ROC curves for a simple threshold filter based on Ab_fa_rep and Ag_fa_rep alone, demonstrating that van der Waals repulsion alone provides substantial discriminative power (nanobody AUC = 0.867/0.858; antibody AUC = 0.893/0.893).

Failed complexes carried substantially higher per-residue fa_rep at the interface than successful ones, for both conventional antibodies and nanobodies (Figure 2E, left and center panels; p < 0.0001). Structural inspection of representative failed cases revealed that individual interface residues can accumulate extreme van der Waals repulsion energies exceeding 600-1,000 REU, indicating severe atomic clashes at the predicted interface (Figure 2C). A comparison of interface interaction properties between representative failed and successful predictions further confirmed that failed cases exhibit fewer stabilizing contacts (hydrogen bonds, salt bridges) despite comparable or higher numbers of total interface contacts (Figure 2D).

We next asked whether van der Waals repulsion alone could serve as a quality filter. ROC analysis using Ab_fa_rep and Ag_fa_rep as threshold classifiers demonstrated substantial discriminative power for both nanobody-antigen (AUC = 0.867/0.858) and antibody-antigen (AUC = 0.893/0.893) complexes (Figure 2E, right panels), eliminating approximately 70% of false-positive predictions while retaining 90% of true-positive cases. However, the remaining 30% of false positives — those with moderate fa_rep values — require a more comprehensive feature set for accurate discrimination, motivating the machine learning approach described in the next section.

### 3. Machine Learning-Based Filter for AF3 Prediction Quality

The threshold-based fa_rep filter described above removed approximately 70% of false positives, but its reliance on only two features limits recall of incorrect predictions that exhibit subtler energetic anomalies. To capture the full discriminative power of the 36-dimensional interface energy feature space, we trained and compared five machine learning classifiers using the top 10 features ranked by Random Forest importance (Figure 2B), as detailed in Methods.

The workflow of the resulting AFilter framework — data collection, AF3 prediction, and machine learning-based quality assessment — is summarized in Figure 3A, with the data sources and partitioning strategy shown in Figure 3B. Among the algorithms evaluated, Random Forest achieved the highest mean AUC for nanobody-antigen classification (AUC ≈ 0.95), followed closely by GBDT (Figure 3C, left). For antibody-antigen classification, Random Forest and GBDT yielded comparable AUC values (Figure 3C, right), though the difference was not statistically significant across the 100 random-seed replicates. Given Random Forest’s comparable performance and greater stability at the current dataset scale, it was selected as the final classifier for both antibody and nanobody filtering (see Methods for selection rationale).

**Figure 3.**
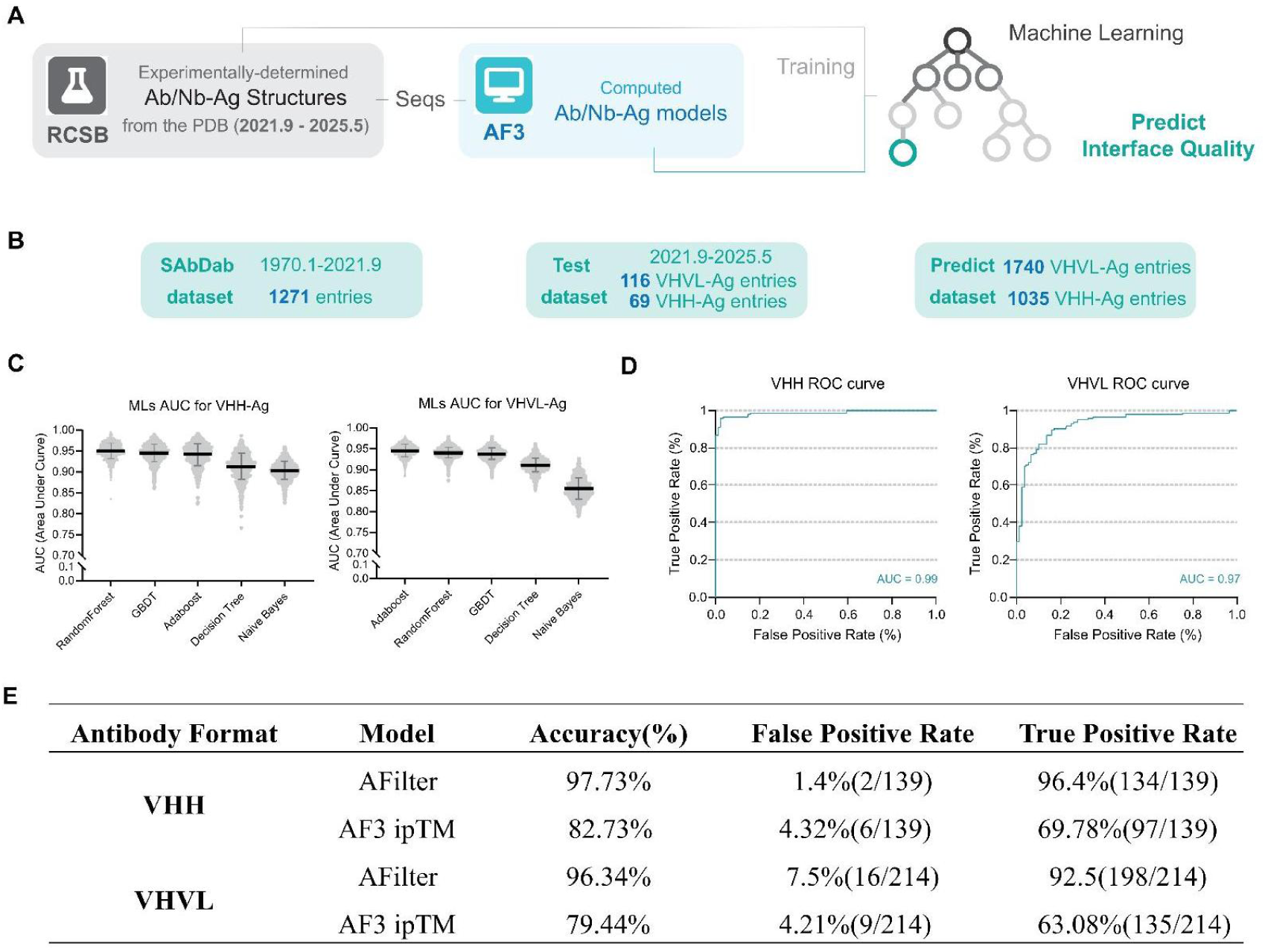
AFilter: Machine Learning-Based Quality Filter for AF3 Predictions. (A) Schematic of the AFilter framework. Experimentally determined antibody/nanobody-antigen structures from the PDB are used to generate AF3 predictions, which are then scored by Rosetta energy features and classified by a trained machine learning model to predict interface quality. (B) Data sources and partitioning. The SAbDab reference dataset (1,271 entries, 1970-2021.9), the temporally held-out test dataset (116 VHVL-Ag and 69 VHH-Ag entries, 2021.9-2025.5), and the AF3 prediction dataset (1,740 VHVL-Ag and 1035 VHH-Ag conformations) are shown. (C) Comparison of classifier AUC values across machine learning algorithms (Random Forest, GBDT, AdaBoost, Decision Tree, Naive Bayes) for VHH-Ag (left) and VHVL-Ag (right) classification tasks. Violin plots show AUC distributions across 100 random-seed replicates. (D) ROC curves for the best-performing AFilter Random Forest models on the held-out validation set. Left: VHH-Ag model (AUC = 0.99); right: VHVL-Ag model (AUC = 0.97). (E) Comparison of AFilter and ipTM-based filtering performance on the validation set. For VHH complexes, AFilter achieved 97.73% accuracy with 96.4% true positive rate (TPR) and 1.4% false positive rate (FPR), compared with ipTM-based classification (threshold = 0.8) at 82.73% accuracy, 69.78% TPR, and 4.32% FPR. For VHVL complexes, AFilter achieved 96.34% accuracy with 92.5% TPR and 7.5% FPR, compared with ipTM at 79.44% accuracy, 63.08% TPR, and 4.21% FPR. ipTM-based classification used the maximum chain-pair ipTM value between antibody and antigen chains extracted from AF3 summary_confidences output.

To prioritize retention of true positives, the classification threshold was tuned to maintain a high true positive rate. The best-performing Random Forest models achieved AUC = 0.99 for VHH-Ag (nanobody-antigen complexes, VHH: nanobody) and AUC = 0.97 for VHVL-Ag (antibody-antigen complexes, VHVL: conventional antibody) on the held-out validation set (Figure 3D). On the validation set, AFilter achieved an overall accuracy of 97.73% for VHH complexes (true positive rate = 96.4%, false positive rate = 1.4%) and 96.34% for VHVL complexes (true positive rate = 92.5%, false positive rate = 7.5%) (Figure 3E). To contextualize these gains, we evaluated AF3’s own internal confidence metric — the interface predicted TM-score (ipTM) — as a baseline binary classifier using the same dataset. Applying a threshold of 0.8, ipTM-based filtering achieved an accuracy of 82.73% for VHH complexes (true positive rate = 69.78%, false positive rate = 4.32%) and 79.44% for VHVL complexes (true positive rate = 63.08%, false positive rate = 4.21%) (Figure 3E). Although ipTM maintained a low false positive rate comparable to AFilter, its substantially lower true positive rate indicates that ipTM discards approximately 30–37% of correctly docked conformations — a critical limitation for epitope mapping, where maximizing recovery of true positives is paramount. AFilter thus achieved a 15–17 percentage-point improvement in accuracy and a 27–30 percentage-point improvement in true positive rate over ipTM-based filtering, while maintaining a comparable or lower false positive rate.

### 4. Enhancing Antibody Epitope Prediction with AFilter

To assess AFilter’s generalizability beyond the training distribution, we evaluated the optimized Random Forest models on an independent test set comprising 12 antibody-antigen and 12 nanobody-antigen complexes experimentally determined between June and November 2025, selected using the same resolution and redundancy criteria applied to the training data (Figure 4A). For each pair, AF3 generated 15 conformations under the standard three-seed protocol.

**Figure 4.**
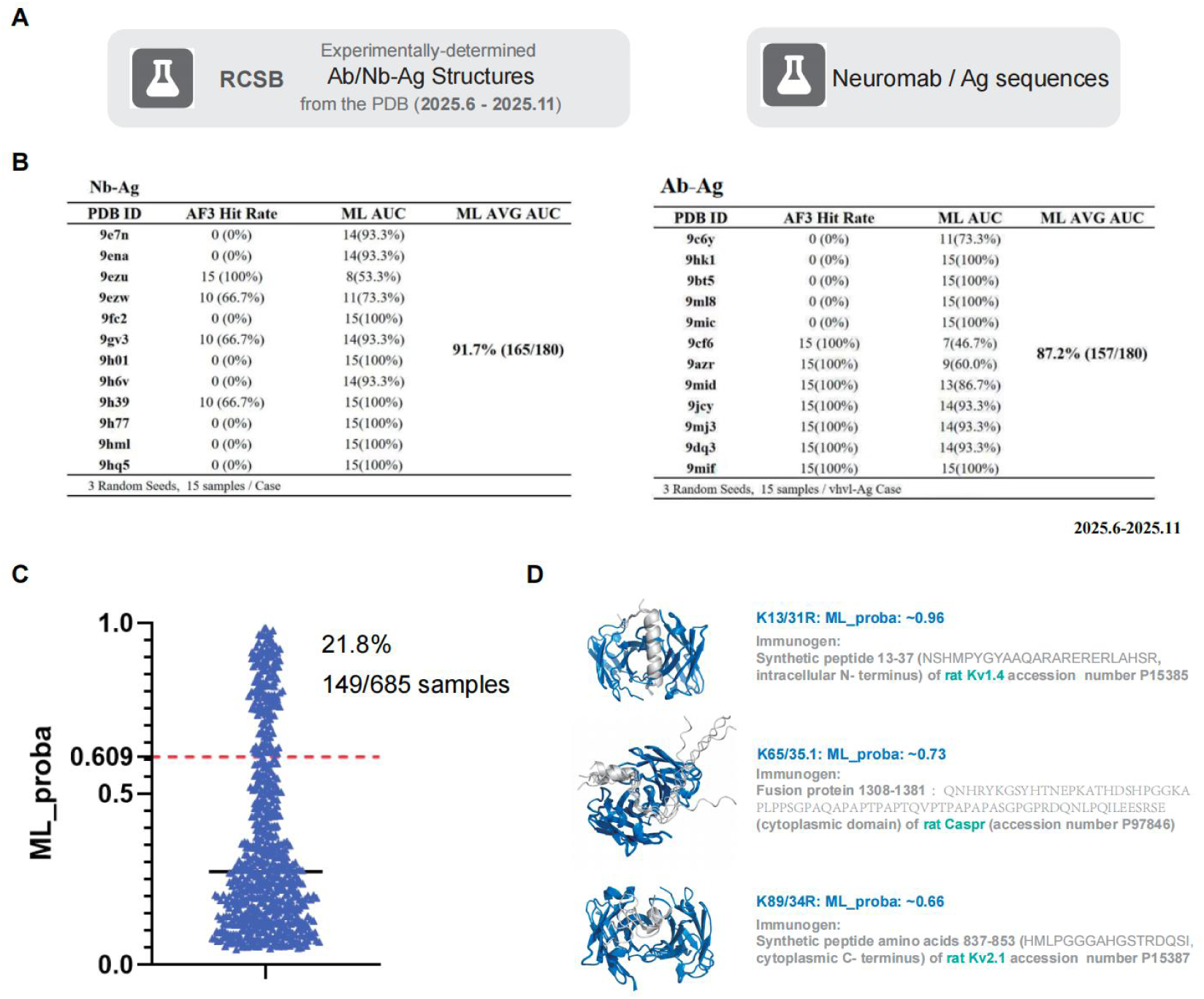
Independent Validation and Practical Application of AFilter. (A) Data sources for independent evaluation. Newly deposited antibody/nanobody-antigen complexes from the PDB (June-November 2025) serve as the external test set, and NeuroMab antibody/antigen sequence pairs serve as the practical application case. (B) Detailed AFilter classification results on the independent test set. Left table: 12 nanobody-antigen complexes with per-PDB AF3 hit rates and AFilter classification accuracy (overall 91.7%, 165/180). Right table: 12 antibody-antigen complexes with per-PDB results (overall 87.2%, 157/180). Three random seeds with 15 samples per case were used. (C) Distribution of AFilter-predicted probabilities (ML_proba) for 685 NeuroMab antibody-antigen prediction samples. The dashed red line indicates the classification threshold (0.609); 21.8% of samples (149/685) were retained as high-confidence predictions above this threshold. (D) Representative NeuroMab application examples showing AFilter-retained high-confidence epitope predictions. Three antibody-antigen pairs are shown with their AFilter probability scores and immunogen descriptions: K13/31R targeting rat Kv1.4 (ML_proba ∼0.96), K65/35.1 targeting rat Caspr (ML_proba ∼0.73), and K89/34R targeting rat Kv2.1 (ML_proba ∼0.66).

On this external test set, AFilter achieved an overall classification accuracy of 91.7% (165/180) for nanobody-antigen pairs and 87.2% (157/180) for antibody-antigen pairs (Figure 4B). Detailed per-complex results revealed that AFilter correctly classified the vast majority of predictions even for complexes where AF3 achieved a 0% hit rate, confirming that AFilter reliably identifies incorrect docking poses. In both cases, the false negative rate remained low, indicating that AFilter seldom discards correctly docked conformations. These results demonstrate that the energy-based filtering strategy generalizes to complexes unseen during both AF3 training and AFilter model development.

To illustrate a practical application, we used AFilter to prioritize candidate epitope predictions for antibodies from the UC Davis/NIH NeuroMab Facility, a public repository of monoclonal antibodies validated for mammalian brain research. Although NeuroMab provides extensive antibody-antigen sequence data, detailed epitope information is largely unavailable for its collection. For each NeuroMab antibody-antigen pair, we generated AF3 structural predictions and applied AFilter to retain only high-confidence conformations. Of 685 total prediction samples, 149 (21.8%) exceeded the AFilter probability threshold of 0.609 and were retained as high-confidence predictions (Figure 4C). Representative examples are shown in Figure 4D, where AFilter assigned high probabilities to structurally plausible predictions for three NeuroMab antibodies targeting neuronal ion channels and cell-adhesion molecules: K13/31R against rat Kv1.4 (ML_proba ∼0.96), K65/35.1 against rat Caspr (ML_proba ∼0.73), and K89/34R against rat Kv2.1 (ML_proba ∼0.66). The computational overhead introduced by the Rosetta energy calculation and AFilter classification steps was minimal, adding less than 5% to the total wall-clock time of the AF3 prediction pipeline.

## Discussion

Our analysis reveals a consistent mechanistic pattern in AF3’s antibody-antigen failures: when the correct binding mode lies outside the model’s accessible diffusion trajectory, AF3 forces the antibody onto an incorrect epitope by remodeling CDR loops — particularly CDR H3 — at the cost of non-physical van der Waals repulsion and backbone strain. This signature is invisible to AF3’s internal confidence metrics but is readily quantifiable by physics-based energy evaluation. A recent independent study corroborates this interpretation, showing that AF3 and related models generate structurally plausible but non-specific antibody-antigen interfaces that cannot be distinguished from correct predictions by confidence scores alone [36]. The phenomenon likely reflects a general limitation of diffusion-based structure generators operating under constrained sampling budgets: with only 15 conformations per pair in our protocol, the probability of encountering the native binding mode is low for challenging targets, and the model defaults to locally optimized decoys [17, 18].

The inadequacy of AF3’s internal confidence metrics for antibody-antigen quality assessment is quantitatively underscored by our ipTM benchmarking results (Figure 3E). The low true positive rate is particularly problematic: ipTM misclassifies approximately one-third of correctly docked conformations as failures, substantially reducing the yield of usable predictions. This finding is consistent with the observation by Smorodina et al. [36] that ipTM fails to discriminate real from shuffled antibody-antigen complexes, and extends it by demonstrating that even within cognate pairs, ipTM cannot reliably identify correct docking poses. AFilter grounds quality assessment in physics-based interface energy evaluation rather than model self-confidence, achieving a 15–17 percentage-point improvement in accuracy and a 27–30 percentage-point improvement in true positive rate over ipTM alone (Figure 3E).

AFilter addresses this quality gap by exploiting the energetic signatures that differentiate native-like from artifactual interfaces. By training a Random Forest classifier on Rosetta-derived interface energy features, our approach filtered out over 90% of false-positive predictions while retaining approximately 90% of true positives on both validation and independent test data. This strategy is conceptually aligned with recent hybrid approaches that combine deep learning-based structure generation with physics-based rescoring — such as ReplicaDock 2.0, which uses Rosetta replica-exchange docking to refine AlphaFold-generated templates [25], and AI-augmented physics-based docking pipelines that rescore AF3-generated ensembles to achieve performance comparable to AF3 itself [26]. AFilter differs from these approaches in that it operates as a post-hoc classifier rather than a refinement engine, making it computationally lightweight (<5% overhead) and directly applicable to any AF3 output without re-docking.

### Positioning AFilter within the Emerging Landscape of Antibody-Antigen Scoring Methods

During 2024-2026, several computational methods have been developed that share the overarching goal of improving the reliability of AI-predicted antibody-antigen complexes, yet each addresses a different aspect of the problem with a distinct algorithmic design. Clarifying these differences is essential for understanding AFilter’s niche and its complementarity with existing tools.

PPIscreenML [37] was developed to distinguish genuinely interacting protein pairs from non-interacting ones by training a Random-Forest classifier on structural features extracted from AlphaFold2-generated models of both positive and negative (decoy) pairs. Its design philosophy is closely related to AFilter’s: both use RF classifiers on AF-output features to perform binary classification. However, the two methods differ in scope and feature space. PPIscreenML addresses the general protein-protein interaction screening problem and uses seven Rosetta-derived whole-complex features (e.g., total energy, shape complementarity) to classify whether two proteins interact at all. AFilter, by contrast, is purpose-built for antibody-antigen complexes and operates on per-residue interface energy decompositions stratified by antibody-side and antigen-side contributions (36 dimensions), enabling it to detect the CDR-specific geometric distortions (e.g., per-residue fa_rep, rama_prepro) that are characteristic of incorrectly docked antibody poses. Moreover, AFilter explicitly evaluates the quality of a predicted docking pose (correct vs. incorrect epitope) rather than predicting whether binding occurs, making it complementary to interaction-screening tools in a sequential pipeline.

AlphaRED / ReplicaDock 2.0 [25] takes the AF-predicted structure as a starting template and applies Rosetta replica-exchange Monte Carlo docking to re-sample the binding landscape, thereby improving Ab-Ag docking success rates by approximately 50% on the AF failure subset. This physics-based refinement approach is powerful but computationally intensive: each complex requires multiple Rosetta trajectory hours, compared with the seconds-scale overhead of AFilter’s energy extraction and classification. The two methods are therefore complementary rather than competing: AFilter can serve as an upstream triage step that identifies which AF3 outputs are worth investing the additional compute of replica-exchange refinement, and which can be confidently accepted or rejected outright. We note that AFilter functions as a post-hoc filter rather than a re-docking engine, and its applicability is broader because it does not require access to the Rosetta docking infrastructure.

The AI-augmented physics-based docking pipeline of Gaudreault et al. [26] uses AI-guided antibody modeling tools (e.g., IgFold, ABodyBuilder2) to generate CDR-diverse ensembles that are then docked and rescored with ClusPro, demonstrating that physics-based docking combined with AI-generated structural diversity can achieve performance on par with AF3 itself. Like AFilter, this approach recognizes the value of physics-based assessment for antibody-antigen interfaces; however, it operates as a complete docking pipeline rather than a lightweight post-processing filter. AFilter’s advantage lies in its minimal computational footprint and its ability to plug directly into existing AF3 workflows without requiring alternative structure-generation tools.

NanoBinder [38] is the method most architecturally similar to AFilter: it trains a Random Forest classifier on Rosetta energy scores to predict nanobody-antigen binding. Both methods share the insight that Rosetta-derived energy features encode sufficient information to discriminate correct from incorrect nanobody-antigen associations. However, NanoBinder is designed to predict whether a given nanobody-antigen pair will bind (a binary binder/non-binder classification), whereas AFilter assesses whether a specific predicted complex structure is geometrically correct (a docking-quality classification). This distinction is critical because a nanobody may genuinely bind its target while AF3 places it at the wrong epitope — a scenario that NanoBinder would miss but AFilter would detect. Furthermore, NanoBinder uses whole-complex Rosetta scores from relaxed structures (e.g., dG_cross, dSASA) and is restricted to nanobody-only complexes, whereas AFilter uses per-residue interface energy decompositions and is trained and reported as two parallel models that explicitly separate conventional antibody-antigen and nanobody-antigen complexes, reflecting the markedly different CDR-loop geometries and interface topologies of these two recognition modes.

AntiConf [39] introduces a composite confidence metric for AF-predicted antibody-antigen complexes by combining pTM and pDockQ2 through a weighted average, achieving improved precision and recall over individual AF confidence scores. AntiConf operates entirely within the AF confidence-score space and requires no external energy calculations, making it extremely fast. However, our results and those of Smorodina et al. [36] demonstrate that AF internal confidence metrics — including derivatives such as pDockQ2 and ipTM — are fundamentally limited in their ability to discriminate correct from incorrect antibody-antigen interfaces, because they reflect model certainty about its own prediction rather than physical realism of the interface. Our direct benchmarking of ipTM as a binary classifier provides quantitative evidence for this limitation, showing that confidence-score-based approaches sacrifice approximately one-third of true positives to maintain a low false positive rate (Figure 3E). AFilter sidesteps this limitation entirely by grounding its assessment in physics-based energy evaluation, which captures non-physical steric clashes and backbone strain that confidence scores cannot detect. The two approaches are complementary: AntiConf can provide a rapid first-pass ranking, while AFilter supplies the physics-based verification layer needed to identify and remove structurally implausible poses that retain high confidence scores.

ABAG-Rank [40] employs a DeepSets-based deep neural network to rank AF3-predicted antibody-antigen models by learning from geometric interface descriptors (pairwise CA distances), AF3 confidence metrics (PAE, ipTM), and ESM2 protein language model embeddings. It processes variable-sized decoy ensembles in a permutation-invariant manner and is trained with a composite regression-plus-ranking loss function. ABAG-Rank substantially outperforms AF3’s internal ranking and the deep-learning baseline DeepRank-Ab in selecting the best model from an ensemble. However, ABAG-Rank is designed as a ranking model — it orders predictions within an ensemble by relative quality — rather than a binary quality filter that classifies individual predictions as correct or incorrect. AFilter addresses a different and complementary task: it provides an absolute accept/reject decision for each predicted structure based on its physical plausibility, independent of the other predictions in the ensemble. This means AFilter can identify when an entire ensemble lacks any acceptable prediction (a scenario in which ranking is uninformative), whereas ABAG-Rank always returns a top-ranked model regardless of absolute quality. Furthermore, ABAG-Rank requires AF3 PAE matrices and ESM2 embeddings as input features, tying it to the AF3 ecosystem, whereas AFilter relies only on the predicted coordinates and their Rosetta energy decomposition.

Finally, Smorodina et al. [36] provide the most comprehensive demonstration to date that AF3, Boltz-2, and Chai-1 all generate structurally plausible antibody-antigen interfaces even for non-cognate (shuffled) pairs, and that internal confidence scores (ipTM) largely fail to discriminate real from shuffled complexes. Their work frames the specificity problem rather than offering a solution. AFilter directly addresses the quality gap identified by Smorodina et al. by detecting the elevated van der Waals repulsion and backbone strain that accompany non-native interfaces. It provides an orthogonal, physics-grounded discrimination signal that is absent from confidence-score-based approaches. We note, however, that our current study evaluates AFilter on cognate antibody-antigen pairs with known experimental structures and has not yet been tested on the non-cognate discrimination task; extending AFilter to the specificity-screening setting described by Smorodina et al. is an important direction for future work.

In summary, AFilter occupies a distinct position within this ecosystem: it is a lightweight, physics-based, binary quality filter that operates at the per-prediction level, requires no re-docking or upstream model modification, and is agnostic to the generative model. Because its input is only a coordinate file and its output is a Rosetta REF2015 energy decomposition, the framework should in principle transfer to AF3-like open-source generators (Boltz-1, Chai-1) and to future diffusion-based docking models without retraining — although this cross-predictor transferability remains to be tested and is a direction for future work.

### Broader Implications and Limitations

A broader issue highlighted by our findings is the tendency of current structure prediction models to “hallucinate” protein-protein interfaces — generating physically reasonable but biologically non-specific complexes even when presented with non-cognate or weakly interacting pairs. This behavior has been attributed to training set bias: the PDB is dominated by genuine complexes, providing few examples of non-interacting pairs from which models could learn to predict unbound conformations [36, 37, 41]. Mischley et al. demonstrated that a dedicated ML classifier (PPIscreenML) trained on AF2-generated models of both interacting and non-interacting protein pairs can substantially improve discrimination, suggesting that explicit inclusion of negative examples in training data may partially alleviate this bias [37]. In the antibody context, such negative examples could consist of antibody-antigen pairs with known non-binding relationships, presented at randomized spatial separations during training. AFilter sidesteps this training-data limitation by operating downstream of structure generation, classifying poses on the basis of their physical realism rather than their assumed binding probability.

Several limitations of the current study should be noted. First, AFilter’s performance depends on the quality of Rosetta energy evaluation, which itself has known limitations for flexible loop regions and may not fully capture solvation effects at antibody-antigen interfaces [24]. Second, our training and test sets, while temporally separated from AF3’s training data, are drawn from the same structural universe (PDB-deposited complexes) and may not fully represent the diversity of therapeutic antibody targets encountered in drug discovery. Third, AFilter is designed to filter, not to correct: it can identify poor predictions but cannot recover the correct binding mode if it is absent from the AF3 sampling ensemble. Increasing sampling depth — by using more seeds or enhanced sampling strategies-remains necessary to improve the probability of generating near-native conformations in the first place.

Looking forward, several directions could extend the utility of this framework. Incorporating Rosetta relaxation prior to energy extraction may further sharpen the energetic contrast between correct and incorrect predictions, potentially improving discrimination in borderline cases. Integrating van der Waals repulsion terms directly into the AF3 loss function during fine-tuning could reduce the generation of steric clashes at source, potentially improving both AF3’s raw success rate and AFilter’s downstream discrimination. Together, these directions could further strengthen the reliability of computational antibody epitope prediction.

## Acknowledgments

We thank Dr Sheldon Xie for providing suggestions on the design and validation of the random forest algorithm.

## Author contributions

X.L. conceived the study, performed the analysis; Y.W. conceived and supervised the study. X.L and Y.W. wrote the paper.

